# Social stress worsens colitis through β-adrenergic–driven oxidative stress in intestinal mucosal compartments

**DOI:** 10.1101/2025.07.15.664961

**Authors:** Elisa Caetano-Silva, Miranda Hilt, Ivan Valishev, Casey Lim, Mikaela Kasperek, Akriti Shrestha, Robert McCusker, Heather Armstrong, Brett Loman, Michael Bailey, Jacob M. Allen

## Abstract

Psychological stress is a known risk factor for inflammatory bowel disease (IBD), but the mechanisms linking stress to worsened disease remain unclear. Because distinct stress paradigms activate different neuroimmune circuits, it is critical to investigate model-specific effects. We examined how social stress primes the gut for heightened inflammation and whether this is mediated by specific neuroendocrine pathways, including α2-/β-adrenergic (sympathetic) or glucocorticoid/ corticotropin-releasing hormone receptor (CRHR1) (HPA axis) signaling. Mice were exposed to social disruption (SDR) stress and pre-treated with pharmacological antagonists targeting α2-adrenergic receptors (idazoxan), β-adrenergic receptor (β-AR) (propranolol), glucocorticoid receptor (mifepristone), or CRHR1 (antalarmin). Intestinal epithelial cell (IEC) gene expression and microbiota composition were assessed following SDR. To determine disease impact, SDR was combined with either *Citrobacter rodentium* infection or dextran sulfate sodium (DSS)-induced colitis, with interventions including the β-AR inhibitor propranolol and the NADPH oxidase inhibitor apocynin. SDR significantly upregulated expression of Dual oxidase 2 *(Duox2)*, Dual oxidase maturation factor 2 *(Duoxa2)*, and inducible nitric oxide synthase 2 *(Nos2)* in IECs (2- to 8- fold, *p* < 0.0001), effects reversed by β-AR blockade but not α2-adrenergic, CRH, or glucocorticoid inhibition. SDR also induced microbial dysbiosis, characterized by reduced alpha-diversity and compositional shifts, which was rescued by propranolol. Stress exacerbated disease severity in both infectious (*C. rodentium*) and chemically induced (DSS) colitis, amplifying colonic expression of *Duox2*, *Nos2*, and *Ccl2,* especially. Apocynin mitigated stress-induced ROS/RNS production and body weight loss even prior to colitis onset, reduced colonic gene expression of key oxidative enzymes, and alleviated both chemically and infectious colitis severity. These findings provide strong evidence that social stress sensitizes the gut to inflammation through β-adrenergic and NADPH oxidase–driven oxidative stress, highlighting potential therapeutic targets for mitigating stress-exacerbated IBD.

**Highlights:** Social disruption (SDR) and restraint stress (RST) activate distinct neuroendocrine pathways, with SDR driving epithelial ROS/RNS pathways via β-adrenergic signaling.

β-adrenergic blockade prevents SDR-induced epithelial priming, microbial dysbiosis, and colitis exacerbation.

NADPH oxidase inhibition with apocynin mitigates stress-induced oxidative stress and disease severity across different colitis models.

Findings identify β-adrenergic and redox pathways as therapeutic targets for stress-exacerbated IBD.

## 1. Introduction

Inflammatory bowel diseases (IBD), including Crohn’s disease and ulcerative colitis, have increased in prevalence in recent decades and represent a significant global health burden (Agrawal & Jess, 2022; Caron et al., 2024). IBD pathogenesis is multifactorial, arising from complex gene- environment interactions, with growing evidence implicating disrupted brain-gut-microbiota communication as a central contributor (Craig et al., 2022). Among environmental risk factors, psychological stress has emerged as a robust and clinically relevant trigger, with perceived stress, trauma, and adverse life events linked to increased disease onset, relapse, and symptom severity (Bitton et al., 2008; Black et al., 2022).

Physiological responses to stress involve activation of two key neuroendocrine pathways: the sympathetic nervous system (SNS), which rapidly releases catecholamines, and the hypothalamic- pituitary-adrenal (HPA) axis, which promotes glucocorticoid secretion (Godoy et al., 2018). Both pathways influence intestinal physiology, but emerging data suggest they may play distinct roles depending on the nature of the stressor. For example, recent work by Schneider et al. (2023) implicates glucocorticoids as primary mediators of stress-induced colitis, yet the generalizability of this mechanism across stress paradigms remains unclear. Social stress models, such as social disruption (SDR), may preferentially engage SNS signaling and provoke unique host responses that differ from those elicited by restraint or early-life stress (Hanke et al., 2012; Lauten et al., 2025; Wohleb et al., 2011; Yadav et al., 2023). Thus, model-specific interrogation of stress signaling is essential to define the biological mechanisms linking stress to IBD.

Intestinal epithelial cells (IECs), which form the frontline of host-microbiota interaction, are increasingly recognized as central players in IBD pathogenesis (Malik et al., 2023). Beyond acting as a physical barrier, IECs secrete cytokines, antimicrobial peptides, and reactive oxygen and nitrogen species (ROS/RNS) to maintain mucosal homeostasis (Castrillón-Betancur et al., 2023; Soderholm & Pedicord, 2019). ROS/RNS generated by NADPH oxidases—particularly DUOX2—and inducible nitric oxide synthase (iNOS), help defend against pathogens but can also disrupt the mucosal barrier when dysregulated. Indeed, DUOX2 and iNOS are consistently elevated in IBD and correlate with disease severity in human cohorts (Burgueño et al., 2021; Haberman et al., 2014; Urbauer et al., 2024).

Our previous work shows that SDR stress upregulates ROS- and RNS-generating pathways in IECs, sensitizing the epithelium to microbial insults (Allen et al., 2022). Yet, the neuroendocrine mechanisms that elevate epithelial ROS tone and how they influence susceptibility to IBD remain poorly understood. In this study, we investigated the role of SNS-driven versus HPA-driven signaling in mediating social stress-induced modifications to mucosal compartments. Using pharmacologic blockade during SDR stress, we identified β-adrenergic signaling as a key driver of DUOX2 and NOS2 upregulation in IECs. Targeting this pathway in both infectious and chemically induced colitis models revealed that inhibiting β-adrenergic receptors (β-AR) or NADPH oxidases—thereby limiting DUOX2- and iNOS-mediated ROS and RNS production, respectively—provides effective protection against stress-aggravated colitis. These findings suggest that stress-induced colitis susceptibility may arise through distinct hormonal mechanisms depending on the stress context, with catecholaminergic and ROS pathways playing a dominant role in the SDR model.

## 2. Results

### 2.1. Distinct stress paradigms engage divergent neuroimmune pathways to shape ROS/RNS epithelial response and the gut microbiome

To investigate how psychological stress impacts intestinal epithelial and microbial homeostasis, we utilized two established murine stress models: SDR and restraint stress (RST). SDR is characterized by daily exposure to an aggressive intruder for 2 hours over 6 days, whereas RST involves placing mice in ventilated restrainers for 2 hours per day over 6 days. While prior work by the Thaiss group (Schneider et al., 2023) showed that RST worsens colitis via corticosterone-driven mechanisms, our preliminary findings suggested that SDR engages distinct neuroimmune pathways. We therefore hypothesized that SDR and RST differentially regulate IEC responses through divergent upstream mechanisms.

Transcriptomic profiling of IECs (EPCAM+, CD45-) revealed that SDR significantly upregulated oxidative stress–related genes compared to unstressed controls, including *Duox2*, *Duoxa2*, and *Nos2* (**Fig. 1A**). In contrast, RST suppressed expression of these genes (**Fig. 1B**). These findings suggest opposing effects on mucosal redox signaling and implicate ROS/RNS as central to the epithelial stress response in this model. In addition, SDR elevated colonic tissue levels of epinephrine from 1.11 to 1.78 ng/g (a 61% increase) and norepinephrine from 162.1 to 210.0 ng/mg (a 30% increase) compared to unstressed controls (**Fig. 1C-D**), with no change in corticosterone (**Fig. 1E**), indicating a dominant role for local SNS activation over systemic HPA signaling during SDR.

**Figure 1.**
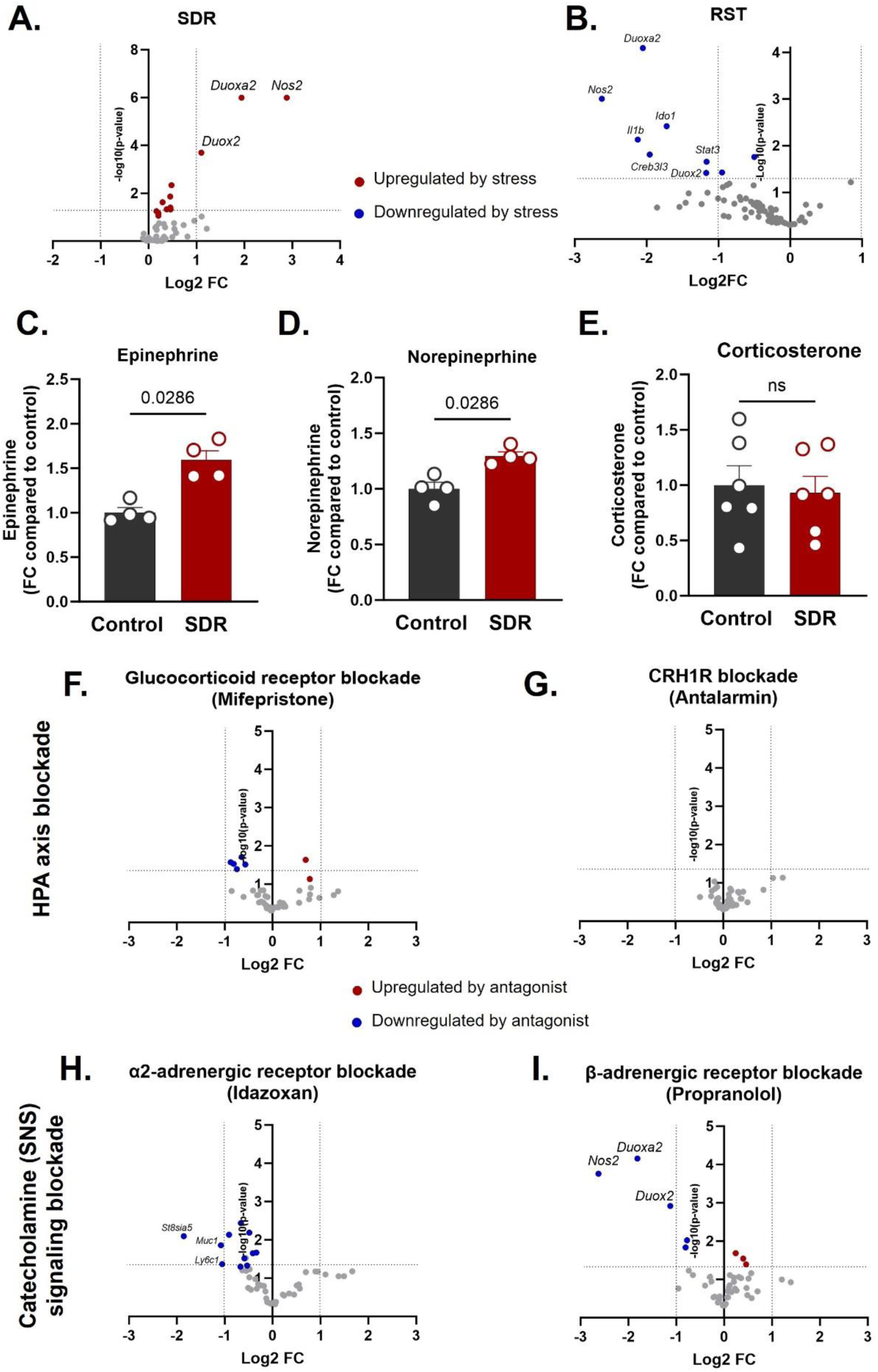
Stress-induced transcriptional changes in intestinal epithelial cells (IECs) are dependent on stressor modality and differentially modulated by blockade of sympathetic nervous system (SNS) versus hypothalamic-pituitary-adrenal (HPA) axis signaling. (A-B) Volcano plots display differential gene expression in IECs following (A) social disruption stress (SDR) or (B) restraint stress (RST), compared to unstressed control animals. Plots show log₂ fold change (x-axis) vs. –log₁₀ p-value (y-axis). Red and blue dots indicate genes significantly upregulated or downregulated by stress, respectively. Horizontal dotted lines indicate significance threshold (*p* < 0.05). (C–E) Colonic levels of (C) epinephrine, **(D)** norepinephrine, and **(E)** corticosterone following SDR, as measured by ELISA. Data are shown as fold change relative to control; p-values from Mann–Whitney test are indicated. **(F–I)** Volcano plots showing IEC gene expression during SDR after pharmacological blockade of key stress hormone receptors: **(F)** glucocorticoid receptor antagonist mifepristone (HPA axis), **(G)** CRHR1 antagonist antalarmin (HPA axis), **(H)** α₂-adrenergic receptor antagonist idazoxan (SNS), and **(I)** β-adrenergic receptor antagonist propranolol (SNS). Each plot shows log₂ fold change vs. –log₁₀ p-value relative to vehicle-treated stressed controls. Red and blue dots represent genes significantly upregulated or downregulated by antagonist treatment, respectively. Horizontal dotted lines indicate significance threshold (*p* < 0.05). n=5-6/group.

To further define the upstream mediators of SDR, mice were pretreated with antagonists targeting glucocorticoid receptors (mifepristone), CRH1R (antalarmin), α2-adrenergic receptors (idazoxan), or β- adrenergic receptors (propranolol) (**Fig. 1F-I**). Only propranolol suppressed SDR-induced expression of *Duox2*, *Duoxa2*, and *Nos2* (**Fig. 1I**), while HPA axis blockade had minimal effects (**Fig. 1F-G**), implicating β-adrenergic signaling as the primary driver of SDR-induced oxidative stress gene transcription. SDR also impaired barrier integrity, as evidenced by serum levels of lipopolysaccharide- binding protein (LBP), a marker of microbial translocation, which was reversed by propranolol (βAR- blocker) (**Suppl. Fig. S1A**).

Given growing evidence that stress alters the gut microbiome (Allen et al., 2022; Beurel, 2024; Karl et al., 2018) and that microbial changes can influence intestinal inflammation, we next investigated which neuroendocrine signaling pathways contribute to microbial dysbiosis. SDR induced significant microbial restructuring, altering Bray-Curtis β-diversity and reducing α-diversity across multiple cohorts (**Suppl. Fig. S1B–F**). Among all antagonists tested, only β-AR blockade with propranolol significantly restored microbial α-diversity (**Suppl. Fig. S1F**), while antagonism of α2-adrenergic, CRH, or glucocorticoid receptors had negligible effects (**Suppl. Fig. S1C-E**), further highlighting the central role of β-adrenergic signaling in stress-induced microbial disruption. Taxonomic composition analysis revealed pronounced shifts in microbial communities following SDR exposure, with numerous taxa significantly upregulated or downregulated compared to non-stressed controls (**Suppl. Fig. S1G**). However, mice subjected to SDR and treated with propranolol exhibited markedly fewer taxonomic changes relative to controls (**Suppl. Fig. S1H**), indicating that blockade of β-adrenergic signaling blunts stress-induced microbial alterations. Together, these findings suggest that SDR drives microbial dysbiosis primarily through β-adrenergic signaling pathways, likely in conjunction with epithelial ROS production.

### 2.2. β-adrenergic receptor blockade mitigates stress-exacerbated infectious colitis

Given the strong effects of β-adrenergic signaling on mucosal ROS and inflammatory pathways, we next tested whether blocking this pathway could prevent stress-induced exacerbation of colitis. We first used *Citrobacter rodentium*, a murine pathogen that recapitulates key features of human enteropathogenic *Escherichia coli* (EPEC) and enterohaemorrhagic *E. coli* (EHEC) infections, as well as modeling some versions of infection-driven IBD (Bouladoux et al., 2017). Mice underwent six consecutive days of SDR stress with or without the β-adrenergic antagonist propranolol. On day 4 of the SDR paradigm, mice were inoculated with *C. rodentium* (**Fig. 2A**), a time point previously identified in preliminary experiments as corresponding to peak stress-induced immune alterations.

**Figure 2.**
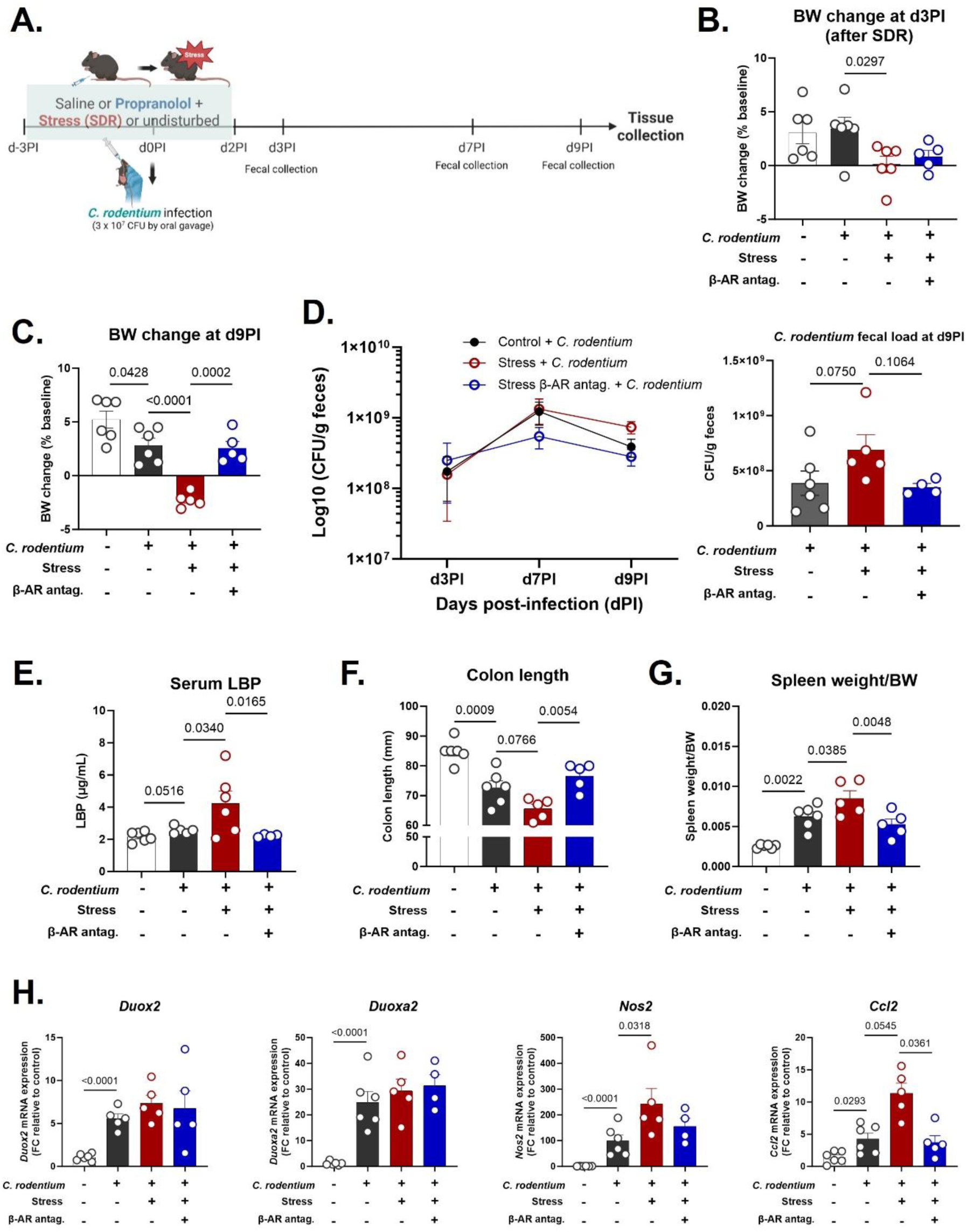
Stress exacerbates *C. rodentium*-induced colitis severity, and β-adrenergic blockade modulates disease progression. **(A)** Timeline for *C. rodentium*-induced colitis + SDR paradigm. dPI = days post-infection. Mice were oral challenged with 3 x 10^7^ CFU of *C. rodentium* at Day 4 of SDR paradigm (d0PI). **(B-C)** Body weight (BW) changes as % of baseline at **(B)** d3PI (after SDR) and **(C)** d9PI. **(D)** *C. rodentium* fecal load at d3PI, d7PI, and d9PI, after bacterial growth in MacConkey agar+ kanamycin for 24h. **(E)** LPS-binding protein (LBP) in serum. **(F)** Colon length; **(G)** Spleen weight to body weight (BW) ratio. **(H)** Fold change gene expression of colonic ROS/RNS-related enzymes (*Duox2, Duoxa2, Nos2*) and chemokine *Ccl2*, relative to control uninfected group. Values were log-transformed prior to analysis. One-way ANOVA was used, followed by post hoc comparisons for selected pairs: Control *C. rodentium* vs. Stress *C. rodentium* and Stress *C. rodentium* vs. Stress Propranolol *C. rodentium*. An unpaired t-test compared Control uninfected and Control *C. rodentium*. Data are presented as mean ± SEM, with *p* < 0.05 considered statistically significant. n=4-6/group.

By d3PI, three days post-infection and before *C. rodentium*-induced weight loss began, stressed mice had already gained less weight than non-stressed controls (*p* < 0.05; **Fig. 2B**), indicating early effects of stress on body weight (BW) regulation. By d9PI, infection decreased BW, but stressed, infected mice exhibited significantly greater BW loss compared to infected-only controls (*p* < 0.0001). Propranolol effectively rescued this weight loss (*p* < 0.001; **Fig. 2C**). Notably, these changes occurred without differences in food intake across groups (**Suppl. Fig. S2A**), suggesting a metabolic effect of stress rather than reduced consumption.

Fecal *C. rodentium* load tended to be higher in stressed animals by d9PI (*p* < 0.10), while propranolol-treated mice showed lower pathogen load across the course of infection (**Fig. 2D**). To evaluate epithelial barrier integrity, we quantified serum levels of lipopolysaccharide-binding protein (LBP), a marker of microbial translocation. *C. rodentium* infection showed a trend towards elevated circulating LBP levels, an effect that was further amplified by stress, yet reversed by propranolol treatment (*p* < 0.05; **Fig. 2E**).

Stress also worsened traditional colitis readouts induced by *C. rodentium*: colon length was significantly reduced in stressed, infected mice compared to unstressed counterparts, and this was ameliorated by β-AR blockade (**Fig. 2F**). Splenomegaly, a marker of systemic immune activation, was elevated in *C. rodentium* infected mice and further so in *C. rodentium* infection plus stress mice, which was reversed by β-AR blockade (**Fig. 2G**). *C. rodentium* infection triggered robust upregulation of oxidative stress-related genes: *Duox2* (8-fold), *Duoxa2* (25-fold), and *Nos2* (98-fold) compared to uninfected controls (*p* < 0.001). Stress further increased *Nos2* expression in infected mice by a further ∼3- fold (*p* < 0.05). Surprisingly, *Duox2* and *Duoxa2* were not significantly impacted by stress or propranolol during *C. rodentium* infection (**Fig. 2H**). Nevertheless, *Ccl2*, a chemokine driving monocyte recruitment, was increased 3-fold by infection and up to 11-fold by stress, with propranolol restoring expression to baseline (*p* < 0.05; **Fig. 2H**).

### 2.3. β-AR signaling also drives stress-aggravated DSS-induced colitis

We next asked whether β-adrenergic signaling contributes to stress-induced worsening of chemically-induced colitis. In a separate cohort, mice underwent SDR stress followed by induction of colitis with 2% DSS in drinking water for 5 days (**Fig. 3A**). By Day 7 (before DSS administration), stressed mice had already gained significantly less weight than controls (**Fig. 3B**). During DSS-induced colitis (Day 12), stressed mice showed significantly lower weight gain than non-stressed DSS-treated controls (*p* < 0.05), indicating that stress exacerbated disease-associated weight loss (**Fig. 3C**). However, by the end of the experiment (Day 15), this difference was no longer apparent, possibly due to the robust BW reduction induced by DSS across all groups (**Fig. 3D**). Treatments did not result in consistent reductions in food intake (**Suppl. Fig. S2B**). While propranolol treatment did not prevent DSS-induced weight loss or mitigate the transient exacerbation caused by prior stress, several other clinical markers of disease were significantly responsive to β-AR blockade. Disease Activity Index (DAI), which includes BW loss, stool consistency and blood in stool/rectal bleeding, was significantly elevated in stress + DSS mice by Day 10, compared to control mice (**Fig 4E**), while addition of propranolol displayed reduced DAI beginning at day 13 compared to DSS alone and stress + DSS mice (**Fig. 4E**). By Day 14, stress + DSS mice exhibited moderate to severe colitis (DAI scores of 5–7), whereas their propranolol-treated counterparts showed markedly lower DAI scores (≤4), indicating attenuated disease severity (**Fig. 4E**). Colon shortening caused by DSS was worsened by stress, an effect that was significantly attenuated with propranolol treatment (*p* < 0.05; **Fig. 4F**).

**Figure 3.**
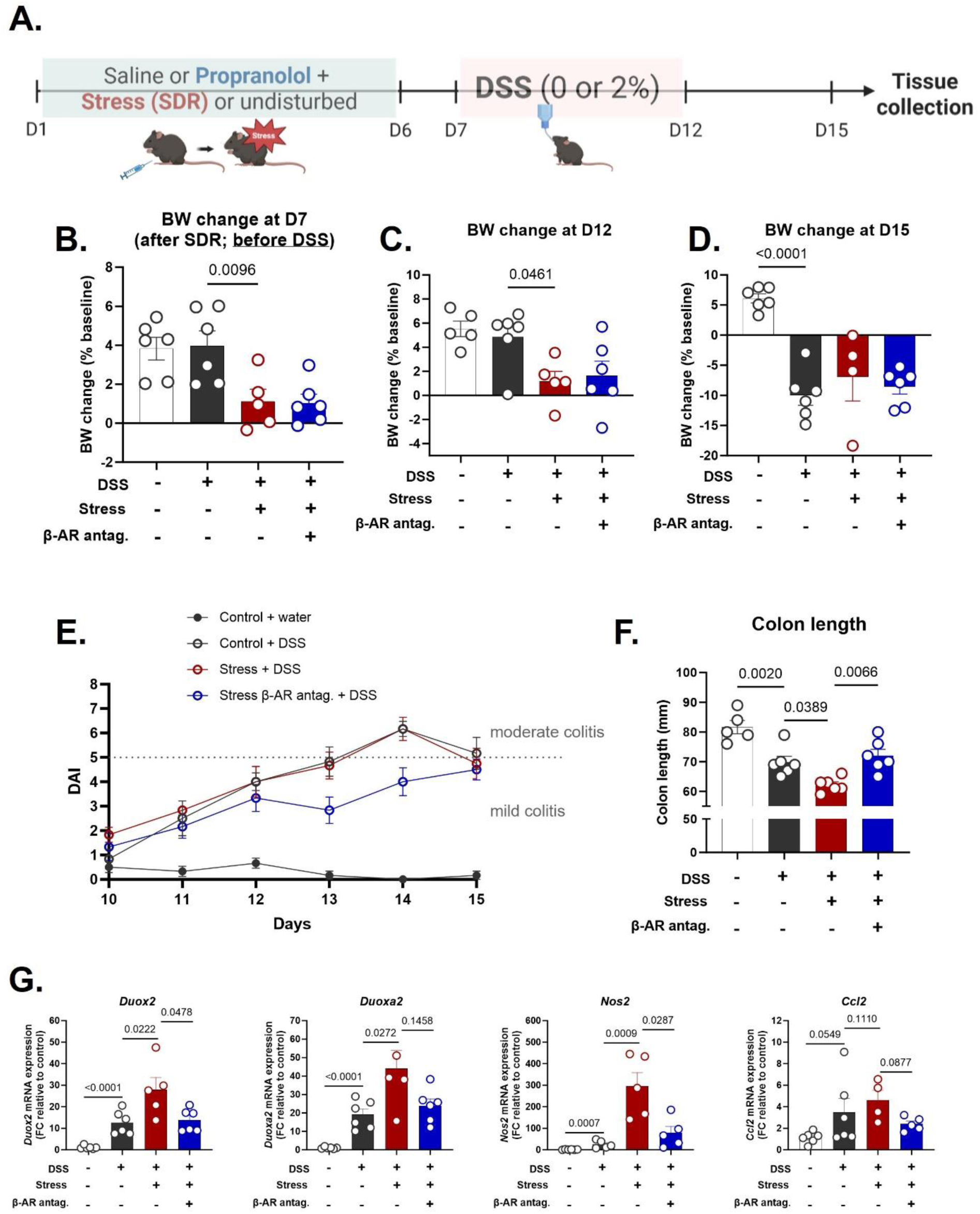
Stress exacerbates DSS-induced colitis severity, and β-adrenergic blockade modulates disease progression. **(A)** Timeline for DSS-induced colitis + SDR paradigm. **(B-D)** Body weight (BW) changes as % of baseline at **(B)** Day 7 (after SDR and prior to DSS administration), **(C)** Day 12, and **(D)** Day 15. **(E)** Disease Activity Index (DAI) on Day 10-15 (3-8 days post-DSS initiation). DAI was determined by scoring weight loss (0 = none, 1 = 1-5%, 2 = 5-10%, 3 = >10%), stool consistency (0 = normal, 1 = soft, 2 = loose, 3 = watery diarrhea), and blood in stool using Hemoccult test (0 = none, 1 = positive, 2 = positive with traces of blood, 3 = positive with gross bleeding). The sum of these scores determined the severity of colitis: 1-4 = mild, 5-7 = moderate, and 8-9 = severe colitis. **(F)** Colon length. **(G)** Fold change gene expression of colonic ROS/RNS-related enzymes (*Duox2, Duoxa2, Nos2*) and chemokine *Ccl2*, relative to control group. Values were log-transformed prior to analysis. One-way ANOVA was used, followed by post hoc comparisons for selected pairs: Control DSS vs. Stress DSS and Stress DSS vs. Stress Propranolol DSS. Data are presented as mean ± SEM, with *p* < 0.05 considered statistically significant. n=4-6/group.

**Figure 4.**
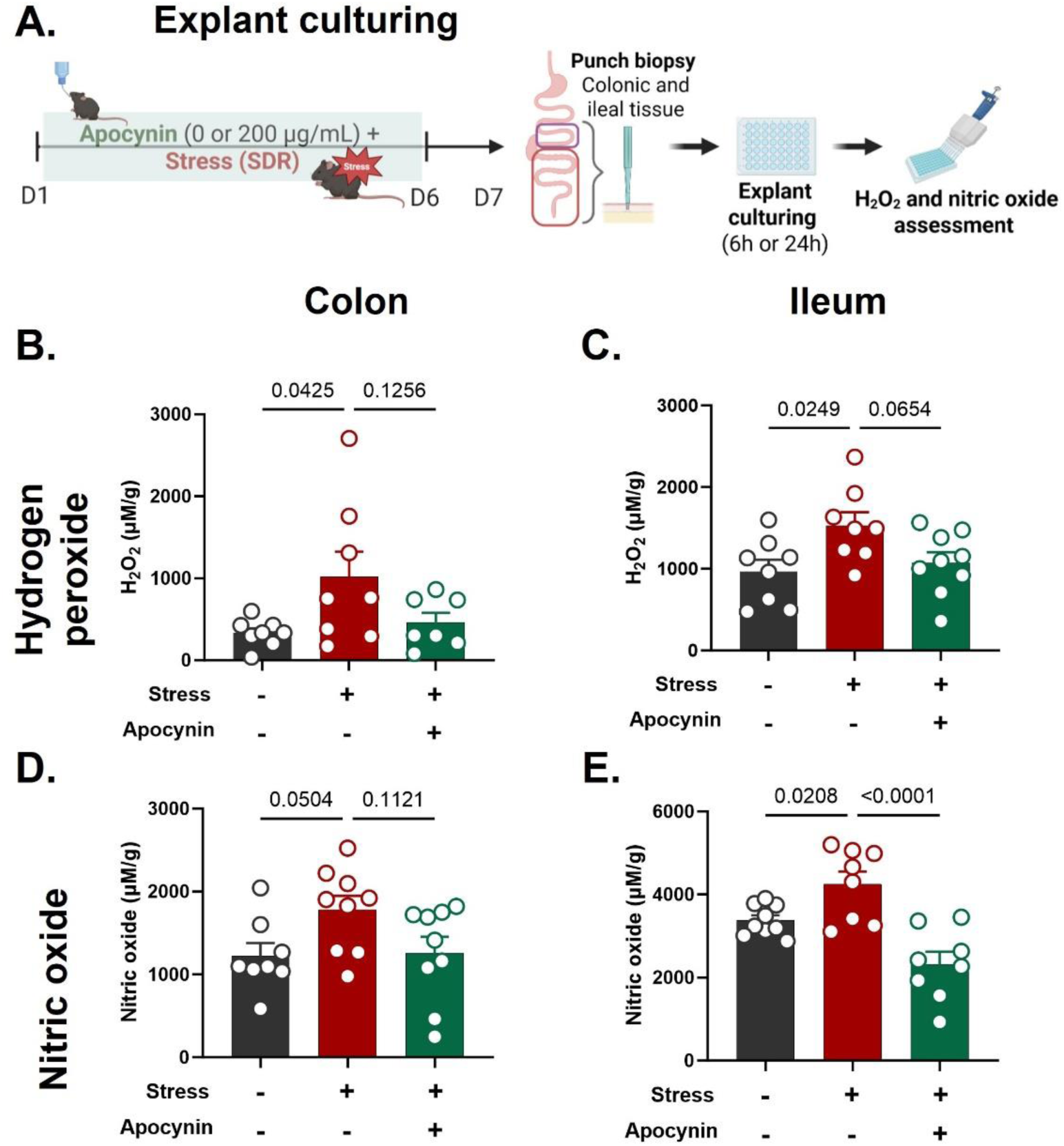
Stress enhances ROS/RNS production in intestinal tissue, which is mitigated by NADPH oxidase inhibition with apocynin in *ex vivo* explant culture. (A) Schematic experimental design. After the 6-day SDR paradigm, **(B and D)** colonic and **(C and E)** ileal biopsies were cultured explant in DMEM/F12-supplemented media using a sponge-based assay. The supernatants were collected for **(B-C)** hydrogen peroxide analysis using the Amplex UltraRed assay at 6 hours, and for **(D-E)** nitric oxide analysis using the Griess Reagent assay at 24 hours. One-way ANOVA was used, followed by post hoc multiple comparisons. Data are presented as mean ± SEM, with *p* < 0.05 considered statistically significant. n=7-9/group.

DSS increased *Duox2*, *Duoxa2*, and *Nos2* expression by 8- to 32-fold, and stress further amplified this ROS/RNS response by 2- to 16-fold over DSS alone. Propranolol significantly reduced this stress-induced enhancement (*p* < 0.05; **Fig. 3G**), providing further support that β-AR signaling mediates social stress driven mucosal ROS signaling.

### 2.4. Apocynin attenuates stress-induced exacerbation of intestinal ROS and RNS

To determine whether ROS signaling plays a direct role in stress-induced colitis severity, we targeted NADPH oxidases, the primary enzymes responsible for mucosal ROS production. We used apocynin, a well-characterized inhibitor that blocks the assembly of the NADPH oxidase complex and prevents downstream ROS generation, including hydrogen peroxide (H₂O₂). First, mice received apocynin via drinking water during the SDR stress paradigm to establish effects on gut-specific ROS levels (**Fig. 4A**). In *ex vivo* explant cultures, stress increased H₂O₂ and nitric oxide (NO) levels in both colon and ileum, while apocynin attenuated these effects, most notably in the ileum (**Fig. 4B–E**). This regional difference suggests that apocynin more potently blunts stress-induced ROS and RNS signaling in the small intestine, potentially reflecting site-specific sensitivity or differences in redox regulation along the gut. Although apocynin does not directly inhibit iNOS, the reduction in NO levels may reflect an upstream role for ROS in facilitating iNOS expression or activity, highlighting potential redox crosstalk between ROS and RNS pathways.

### 2.5. NADPH oxidase inhibition limits social stress-exacerbated colitis

In light of our findings that apocynin attenuates stress-induced exacerbation of gut ROS production, we next asked if these effects could also limit disease outcomes associated with heightened ROS activity. To test this, we treated stressed mice with apocynin in both *C. rodentium* and DSS colitis models (**Fig. 5A, B**). In both models, apocynin rescued stress-induced BW loss, with higher BW already evident immediately post-stress, even before colitis onset (**Fig. C-E**). Apocynin treatment also alleviated colitis severity as evidenced by improved stool consistency scores (**Fig. 5F)** and increased colon length (**Fig. 5G-H**) compared to stress-colitis controls. Consistent with these phenotypic improvements, apocynin significantly suppressed key stress-amplified inflammatory and oxidative transcripts, reducing *Nos2* and *Ccl2* expression in the infection model (**Fig. 5I**) and downregulating *Duox2*, *Duoxa2*, and *Nos2* in DSS model (**Fig. 5J**), compared to *C.rodentium*+stress and DSS-stress controls, respectively. These transcriptional changes align with apocynin’s effects on explant ROS/RNS production, reinforcing its impact on stress-induced redox signaling. Given our previous findings that *Ccl2* expression is essential for multiple stress-induced changes in infection-driven colitis, including increased colonic macrophage accumulation, heightened inflammatory gene expression, and greater bacterial translocation to the spleen (Mackos et al., 2016), these results support the notion that stress-induced ROS signaling primes the gut for heightened immune infiltration and worsened disease severity. Together, these results demonstrate that NADPH oxidase activity is a critical amplifier of stress-induced gut dysfunction, and that its inhibition reverses both oxidative stress priming and downstream colitis severity.

**Figure 5.**
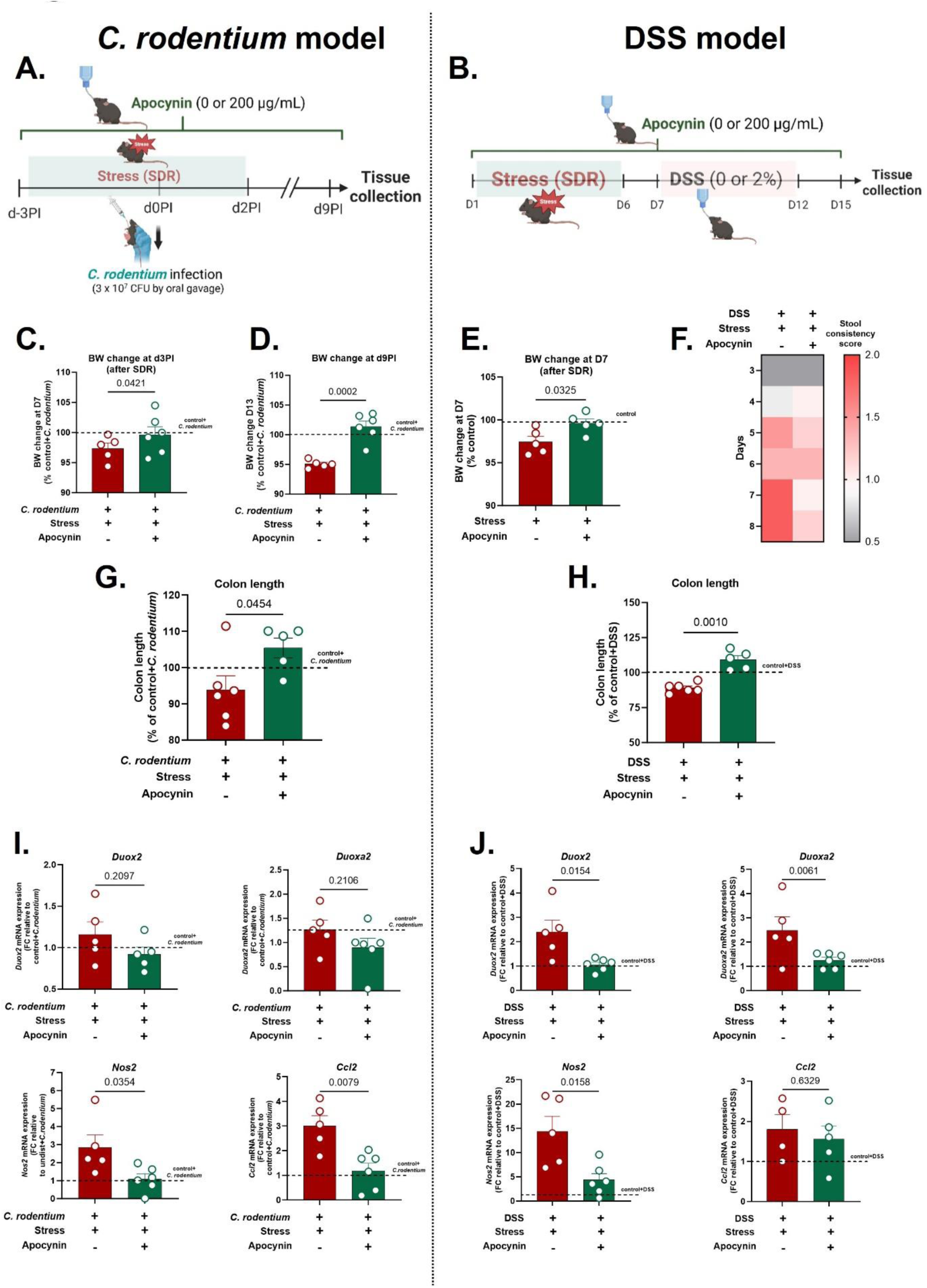
Inhibition of NADPH oxidases by apocynin alleviates stress-induced exacerbation of *C. rodentium*- and DSS-induced colitis. (A-B) Timeline for *C. rodentium*- and DSS-induced colitis, respectively, combined with SDR paradigm. dPI = days post-infection. **(C-E)** Body weight (BW) change at **(C)** d2PI and **(D)** d9PI relative to infected unstressed mice, and at **(E)** D7 relative to water-treated unstressed mice. **(G-H)** Colon length as a percentage of **(G)** infected and **(H)** DSS-treated unstressed mice. **(I-J)** Fold change gene expression of colonic ROS/RNS-related enzymes (*Duox2, Duoxa2, Nos2*) and chemokine *Ccl2*, compared to **(I)** infected and **(J)** DSS-treated unstressed controls. Values were log- transformed prior to analysis and analyzed by t-test. Data are presented as mean ± SEM; *p* < 0.05 was considered statistically significant. n = 5–6/group.

## 3. Discussion

Psychological stress is a known risk factor for human IBD relapse and symptom severity (Black et al., 2022) and exacerbates colitis in preclinical models (Chen et al., 2021; Gao et al., 2018; Mackos et al., 2016), but the underlying mechanisms remain poorly understood. Different stress paradigms appear to engage distinct neuroendocrine circuits, making it critical to define model-specific mechanisms. In this study, social disruption (SDR) and restraint stress (RST) elicited divergent transcriptional responses in intestinal epithelial cells (IECs), consistent with differences in local catecholamine and glucocorticoid levels. These findings highlight that stressor-specific activation of sympathetic or HPA axis signaling differentially shapes mucosal outcomes.

Building on our previous work showing that social stress upregulates ROS- and RNS-generating pathways in IECs, priming the gut for exaggerated inflammatory responses (Allen et al., 2022), we now identify β-adrenergic receptor (β-AR) signaling and oxidative stress as central mediators of this response. β-AR blockade reversed stress-induced transcriptional changes in IECs, including upregulation of *Duox2*, *Duoxa2*, and *Nos2*, and mitigated colitis severity. These effects were not seen with glucocorticoid or CRHR1 antagonism, highlighting a predominant role for catecholaminergic signaling in epithelial priming, in contrast to restraint stress models where glucocorticoids dominate (Schneider et al., 2023; Tena-Garitaonaindia et al., 2022; Zheng et al., 2013).

Intestinal homeostasis relies on continuous crosstalk between the epithelium and the gut microbiota (Burgueño et al., 2019). Consistent with our IEC findings, social stress altered microbial composition, in line with reports from both human (Almand et al., 2022; Delgadillo et al., 2025) and animal studies (Allen et al., 2022; Geng et al., 2020; Shevchenko et al., 2023; Yadav et al., 2023). Notably, only β-AR blockade reversed these shifts, suggesting a key role for catecholaminergic signaling in shaping the microbiota. Elevated intestinal norepinephrine has been linked to stress-induced microbial changes (Yadav et al., 2023), supporting our findings that catecholamines—not glucocorticoids—drive these effects.

Microbial shifts have been strongly implicated in colitis pathogenesis (Wang et al., 2024), and the stress-induced changes here increase colitis susceptibility. Microbial dynamics may also influence the observed ROS/RNS activation in IECs. While nitric oxide (NO) is a key antimicrobial molecule, some pathogens exploit NO derivatives as alternative electron acceptors, to thrive in inflamed, hypoxic environments (Henard & Vázquez-Torres, 2011). These findings suggest that stress promotes a permissive environment for pathogen expansion by disrupting the microbiome and dysregulating host ROS/RNS responses, thereby amplifying colitis susceptibility. Further studies are needed to confirm the contribution of these microbial and host-derived mechanisms to stress-exacerbated disease.

Our findings highlight stress-induced ROS signaling as a key driver of colitis susceptibility. DUOX2, previously identified as a mediator of stress-induced epithelial changes (Allen et al., 2022), regulates mucosal ROS production to modulate host–microbiota interactions but can also drive tissue damage when dysregulated (Sommer & Bäckhed, 2015). Previous studies have demonstrated that intestinal epithelial DUOX2 deficiency protects against colitis, as disruptions in the reciprocal interactions between DUOX2 and the microbiome contribute to disease progression (Castrillón-Betancur et al., 2023). As DUOX2 can both influence and be regulated by microbial signals (Sommer & Bäckhed, 2015), our findings suggest that stress may disrupt this feedback loop, amplifying ROS signaling and microbial imbalance in a way that contributes to disease risk.

We observed that prior stress exposure exacerbated colitis severity in both *Citrobacter rodentium* and DSS-induced colitis models, further amplifying gene expression of ROS-RNS-related enzymes (*Duox2, Duoxa2*, and *Nos2*) and the chemokine *Ccl2*, a key mediator of immune cell recruitment. This aligns with our previous work showing that CCL2-mediated monocyte recruitment is a critical mediator of stress-exacerbated colitis (Mackos et al., 2016). Notably, stress alone does not alter *Ccl2* levels, suggesting that a secondary hit, such as *C. rodentium* infection, can amplify inflammation and immune cell recruitment in a system already primed with elevated ROS levels.

Together, these data suggest that catecholamine signaling drives stress-induced shifts in IECs and microbiota that sensitize the gut to inflammatory insults—though direct effects of propranolol on colitis mechanisms cannot be excluded. While previous studies have demonstrated that propranolol inhibits stress-induced monocyte accumulation in bone marrow, circulation, spleen, and brain that corresponded with reduced pro-inflammatory cytokine production in these tissues (Hanke et al., 2012; Wohleb et al., 2013), to our knowledge, this is the first evidence suggesting such a correlation in the gut.

To further examine the role of ROS in stress-induced colitis exacerbation, we employed the NADPH oxidase inhibitor Apocynin, which effectively mitigated stress-driven weight loss, stool consistency changes, and transcriptional markers of immune activation. Apocynin reduced body weight loss and ROS/RNS production in the gut even before colitis induction, suggesting that oxidative stress contributes to the early inflammatory priming. In the DSS model, apocynin downregulated stress- exacerbated colonic expression of *Duox2, Duoxa2*, and *Nos2*, highlighting NADPH oxidases as key mediators of stress-driven ROS signaling. These findings reinforce the idea that social stress creates a pro- inflammatory gut environment through ROS signaling, predisposing the host to worsened disease outcomes. Given that excessive ROS production from hyperactivated signaling pathways or feedback loops can drive oxidative damage and exacerbate IBD, inhibiting NADPH oxidases may have therapeutic potential. However, both overresponsiveness and suppression NADPH oxidases have been implicated in intestinal inflammatory diseases (Stenke et al., 2019), highlighting the need for future studies to explore different doses of apocynin or other NADPH oxidase inhibitors to better define their role in mitigating stress-induced colitis exacerbation.

In summary, psychosocial stress, particularly social disruption stress which involves strong catecholaminergic activation, enhances ROS signaling, triggers immune activation, and increases colitis severity through β-adrenergic receptor pathways. While various stress paradigms can impact gut physiology, social disruption stress uniquely mimics psychosocial stressors that heavily engage sympathetic outputs, including post-traumatic stress disorders in humans (Lauten et al., 2025; Lauten et al., 2024). Our findings show that both β-AR blockade and NADPH oxidase inhibition effectively reverse stress-induced gut dysfunction, revealing catecholaminergic signaling as a key mechanism linking stress to IBD predisposition and identifying actionable targets for mitigating stress-exacerbated inflammation in IBD.

## 4. Material and methods

### 4.1. Animals

Adult C57BL/6 mice (Charles Rivers Laboratories, Wilmington, MA) 6-8 weeks of age were used for experiments. Mice were given *ad libitum* access to water and standard chow for the duration of experiments, excluding the daily two-hour stress time, during which chow and water were removed for all animals. A 12-hour light/dark cycle was used in the room housing mice (lights on at 5 am). Separate cohorts were used for each antagonist experiment (n=6). All experimental procedures were conducted at the Animal Care Facility at the University of Illinois at Urbana-Champaign with the approval of the Institutional Animal Care and Use Committee (UIUC-Protocol #23225).

### 4.2. Social disruption stress (SDR)

Male mice were exposed to either a social disruption (SDR) stressor or a home cage control for six consecutive days. All SDR experimental groups included only male mice, as this model relies on social defeat and hierarchy formation, which occur robustly in males but not females. Including females would risk mating or uncontrolled aggression, compromising the stress paradigm.

The SDR stressor used was an aggressive CD-1 retired breeder male mouse (Charles Rivers Laboratories, Wilmington, MA) which was added to a cage with three younger and smaller experimental mice at 4 pm (1 hour prior to lights off, i.e. active cycle) for to 2 hours per day. Social avoidance (aka. defeat) of experimental mice was verified if they displayed the classical defeat posture (forelimbs raised, upright posture against the sides of the cage). Mouse sacrifice followed by tissue collection and cell isolation occurred on day seven after the beginning of the six-day SDR protocol.

### 4.3. Restraint stress model (RST)

Male and female mice were subjected to restraint stress for 2 hours per day across six consecutive days, in parallel with the SDR protocol. Beginning 3 hours after lights on, mice were placed in well-ventilated 50 mL conical restraint tubes that restricted movement but allowed normal respiration. Tubes were appropriately sized to prevent injury and stress-induced hypothermia and were sanitized between uses. Mice were continuously monitored throughout each restraint session, then returned to their home cages immediately afterward. Control animals remained undisturbed in their home cages.

Sacrifice and tissue collection occurred on day seven, approximately 24 hours after the final restraint session.

### 4.4. Stress-hormone pharmacological blockade

For each stress/antagonist experiment, mice were randomized to one of three groups: 1) No stress + vehicle; 2) SDR + vehicle; and 3) SDR + antagonist (n=9/group). Antagonist or corresponding vehicle control was administered via intraperitoneal (i.p.) injection (25-gauge needle) daily for six days right before the beginning of 2-hour SDR exposure. Pharmacological antagonists were used to inhibit the major stress hormone receptors: α2-adrenergic receptor (Idazoxan), β-adrenergic receptor (Propranolol), CRH1 receptor (Antalarmin), and glucocorticoid receptor (RU-486/Mifepristone). All antagonists were purchased from Sigma Aldrich (St. Louis, MO, USA). Propranolol hydrochloride (Cat #P0884) and Idazoxan hydrochloride (Cat #16138) were dissolved in saline solution (0.9% NaCl) and administered at a dosage of 10 and 2 mg/kg, respectively. Antalarmin hydrochloride (Cat #A8727) and RU-486 (Mifepristone; Cat #M8046) were dissolved in 10% Tween 80 and 10% DMSO in sterile PBS and administered at a dosage of 20 mg/kg or 50 mg/kg, respectively.

### 4.5. Intestinal epithelial cells (IEC) (CD45-; EpCAM+) isolation

Colons were removed from mice immediately after sacrifice. Luminal contents were first gently collected and separated for microbiome analysis; the tissues were flushed with cold PBS using a gavage needle. Tissues were then opened longitudinally, thoroughly washed with PBS, and cut into 2–3 mm pieces. The tissue pieces were placed in a 50-ml conical tube containing 20 mL pre-digestion solution (1x HBSS, 5 mM EDTA, 1 mM DTT, 5% FBS with Antibiotic/Antimycotic solution (Sigma Aldrich, St. Louis, MO)) and rotated for 20 min at 37°C. After vortexing for 10 seconds, the tissue homogenate was filtered with a 100 µM mesh filter with the resulting pass through (containing colonic cells) placed on ice. The remaining colon pieces were placed in 20 mL of fresh pre-digestion buffer and rotated again. These steps were repeated 3 consecutive times with pass through stored on ice after each rotation to ensure adequate IEC removal from lamina propria. Next, single cell suspensions were diluted to 10^8^ cells/mL in MACs buffer (0.5% BSA 2 mM EDTA). After a wash step, live cells were incubated with CD45 magnetic beads (20 µL per 10^7^ cells) (Cat# 130-052-301, Miltenyi Biotec, Auburn, CA, USA), for 10 min before being passed through Miltenyi MS columns (Cat #130-042-201) per manufacturer’s instructions. The eluted CD45− cell fraction (10^7^–10^8^ cells) was then incubated with EpCAM+ beads (20 µL per 10^7^ cells) (Cat #130-105-958) for 10 min before again passing through MS columns per manufacturer instructions. Resulting cells (CD45-; EpCAM+) were collected and named IECs (intestinal epithelial cells). Cells were stored at -80 °C until further analysis.

### 4.6. Catecholamine and corticosterone assessment in colon tissue

Colonic levels of epinephrine, norepinephrine, and corticosterone were measured using ELISA kits (Adrenaline: Cat #BA E-5100R, Noradrenaline: Cat #BA E-5200R; Version 14.2; Rocky Mountain Diagnostics, CO, USA) according to the manufacturer’s instructions. Colon segments were flash-frozen in liquid nitrogen immediately after collection and stored at –80°C until processing. Tissues were weighed and homogenized in cold extraction buffer (0.01 N HCl, 1 mM EDTA, 4 mM metabisulfite) using tissue homogenizer. Homogenates were centrifuged at 15,000 × g for 15 minutes at 4°C, and supernatants were used for quantification in both assays, epinephrine and norepinephrine.

Colonic corticosterone levels were determined using an ELISA kit (Cat #ab108821; Abcam, Waltham, MA, USA). Colon segments were weighed and homogenized in 1% Triton-X (1 mL/g tissue) using a bead beater. Homogenates were centrifuged at 14,000 × g for 20 minutes at 4°C, and supernatants were collected for analysis.

All samples were run at least in triplicate, and concentrations were normalized to tissue weight.

Data was expressed as fold-change compared to the unstressed control.

### 4.7. Gene expression by Fluidigm

Colon tissue RNA was isolated using the Zymo RNA extraction kit (Cat# R2062, Zymo Research Corporation, Irvine, CA, USA), followed by cDNA synthesis with a High-Capacity cDNA Reverse Transcription Kit with RNA inhibitor (Cat# 4374967, Thermo Fisher Scientific, Waltham, MA USA). Real-Time PCR Fluidigm analysis (96 × 96 chip) was conducted by the University of Illinois at Urbana- Champaign Functional Genomics Unit of the W.M. Keck Center. Data acquisition was performed using the Fluidigm Real-Time PCR Analysis 3.0.2 software (Fluidigm, San Francisco, CA, USA). Relative expression was determined using the delta-delta cycle threshold method (ddCt) with Undisturbed mice as control, and Eukaryotic elongation factor 2 (*Eef2*) served as the housekeeping gene. Values were log2 transformed before statistical analysis. Table S1 shows the Fw and Rv primers used (Integrated DNA Technologies, Coralville, IA).

### 4.8. 16s rRNA microbiome sequencing analysis

Proximal colon contents were removed from animals’ colons immediately after sacrifice, snap frozen in liquid nitrogen, and stored at -80 °C. For sample preparation, proximal colon contents were incubated for 45 min at 37 °C in lysozyme buffer (22 mg/ml lysozyme, 20 mM TrisHCl, 2 mM EDTA, 1.2% Triton-x, pH 8.0) before homogenization for 150 seconds with 0.1 mm zirconia beads. Next, samples were incubated at 95 °C for 5 min with InhibitEX Buffer, followed by incubation at 70 °C for 10 min with Proteinase K and lysis Buffer AL. QIAamp Fast DNA Stool Mini Kit (Cat # 51604, Qiagen, Hilden Germany) was utilized to extract DNA (∼10 mg) from the homogenized content. All conditions followed manufacturer’s instructions, with slight modifications as previously described by Allen et al. (2022). dsDNA Broad Range Assay Kit was used to quantify DNA with Qubit 2.0 Fluorometer (Life Technologies, Carlsbad, CA). Illumina MiSeq was used to obtain amplified PCR libraries sequencing done from both ends of the 250-nucleotide region of V3-V4 16S rRNA hypervariable region. Amplicon processing and downstream taxonomic assignment using the ribosomal RNA database SILVA v138 was performed using the DADA2 and QIIME 2.0 platforms. EMPeror tool was used to visualize the microbial diversity (β-diversity, Unweighted Unifrac) in 3-dimensional PCoA plots.

### 4.9. NADPH oxidase blockade by Apocynin treatment

To assess the blockade of NADPH oxidases, Apocynin (Cat #178385, Sigma Aldrich, St. Louis, MO, USA) was administered via drinking water (200 µg/mL) alongside the SDR paradigm for *ex vivo* culturing, and also used in both colitis models with the SDR paradigm. A stock solution (40 mg/mL) was prepared in 100% ethanol and subsequently diluted in tap water. Fresh solutions were prepared every 2– 3 days, and fluid intake was monitored to ensure adequate consumption. For the DSS model, Apocynin was diluted in a 2% DSS solution instead of tap water, allowing for simultaneous delivery of both compounds.

### 4.10. Colonic and ileal tissue-derived explant culturing and ROS/RNS assessment

After sacrifice of mice fed Apocynin in drinking water and submitted to 6-day SDR course, colon and ileum tissues were collected and cleaned of residual fat and Peyer’s patches. Contents were gently removed, and the tissues flushed with cold PBS using a gavage needle. After opening the tissues longitudinally with scissors, they were rinsed and put in a tube containing cold 2% FBS HBSS for further processing within an hour. Biopsies were collected from the mid-colon and mid-ileum using a sterile disposable biopsy Tru-Punch 6 mm (Sklar, West Chester, PA, USA). *Ex vivo* biopsy cultures were performed following Armstrong et al. (2023) with modifications. Briefly, biopsies were carefully placed onto sterile SURGIFOAM® absorbable gelatin sponge (10 x 10 x 10 mm cube; Cat# 1974, Ethicon, Inc. Somerville, NJ, USA) positioned in 48-well plates. The sponges were soaked but not submerged in supplemented DMEM/F12 containing 10% FBS. Explant tissues were then incubated at 37 °C 5% CO2. The supernatants were collected at 6 hours for hydrogen peroxide analysis using the Amplex Red assay (Cat #A22188) with Amplex Ultra Red reagent (Cat # A36006; Thermo Fisher Scientific, Eugene, OR, USA), and at 24 hours for nitric oxide analysis using the Griess Reagent assay (Cat #G7921).

#### 4.10.1. C. rodentium induced colitis

*Citrobacter rodentium* strain DBS120 (pCRP1::Tn5) was grown in Difco Lennox broth (LB) and prepared as previously described (Mackos et al., 2013). Mice were infected via oral gavage with 100 µL PBS containing 3 × 10^7^ CFU on Day 4, during the ongoing stress paradigm. Mice were deprived of food and water for 2 h before and after challenge to aid in colonization. Fecal samples were collected at baseline (Day 0) and on Days 3, 7, 9 post-infection (dPI) to assess bacterial colonization after growth on MacConkey agar supplemented with kanamycin (40µg/mL) based on prior work (Mackos et al., 2013). Body weight, food and fluid intake were daily monitored. Mice were euthanized on Day 9 dPI, a time point previously determined as the peak of disease severity in a similar *C. rodentium*-induced colitis protocol using this mouse strain. The entire colon (from the cecum to the anus) was excised and measured using digital calipers as an indicator of disease severity. Spleens were also collected and weighed, and the spleen weight-to-body weight ratio was calculated as an additional indicator of systemic inflammation.

#### 4.10.2. Chemically-induced colitis (DSS)

The methodology for the DSS colitis model and monitoring was adapted from previous work (Caetano-Silva, 2024). Dextran sulfate sodium (DSS; molecular weight 36,000–50,000; MP Biomedicals, Illkirch, France) was administered via drinking water at concentrations of 0% or 2% for five days. Colitis induction with DSS began after the 6-day SDR paradigm was completed. A single DSS batch was used across all experiments to ensure consistency. Daily assessments included body weight, food and water intake, stool consistency, and stool blood presence. Starting on the third day of treatment, stool consistency was evaluated using a modified scoring system based on Wirtz et al. (2017), with the following scale: 0 – Normal; 1 – Soft; 2 – Very soft/wet; 3 – Watery diarrhea. Blood in stool was detected using the Hemoccult test (HemoCue America Beckman Coulter™ Hemoccult™ SENSA™ Fecal Occult Blood Slide Test System) and scored as follows: 0 – Negative; 1 – Positive; 2 – Positive with visible traces of blood in stool; 3 – Positive with gross bleeding. Mice were sacrificed on Day 8 after DSS start, a time point previously determined as the peak of disease severity in a similar DSS-induced colitis protocol using this mouse strain (Cook et al., 2013). As in the *C. rodentium* model, colon length and the spleen weight-to- body weight ratio were assessed as indicators of colitis severity and systemic inflammation.

### 4.11. Statistical Analysis

Data are presented as mean ± SEM. Statistical analyses were performed using GraphPad Prism v8.0.1 (GraphPad Software, San Diego, CA), with statistical significance set at *p* < 0.05. Details of the specific statistical tests used are provided in each figure caption.

Microbiome analyses were conducted using the MicrobiomeAnalyst platform (www.microbiomeanalyst.ca) (Lu et al., 2023). Taxonomic classification was based on the SILVA database. Sequencing data were normalized to the minimum library size and transformed using the centered log-ratio (CLR) method. Multiple comparisons were corrected using the Benjamini-Hochberg false discovery rate (FDR). Beta-diversity was evaluated using Bray–Curtis dissimilarity and visualized via principal coordinates analysis (PCoA); significance was tested using PERMANOVA. Alpha-diversity was measured using the Chao1 index. Differential abundance at the genus level was analyzed using multiple linear regression with covariate adjustment and FDR correction. Taxonomic differences between control and stress groups, with or without propranolol treatment, were visualized using heat tree plots (Foster et al., 2017).

## Data availability

Sequencing data will be deposited in the NCBI SRA upon acceptance.

## Supplementary material

**Table S1.**
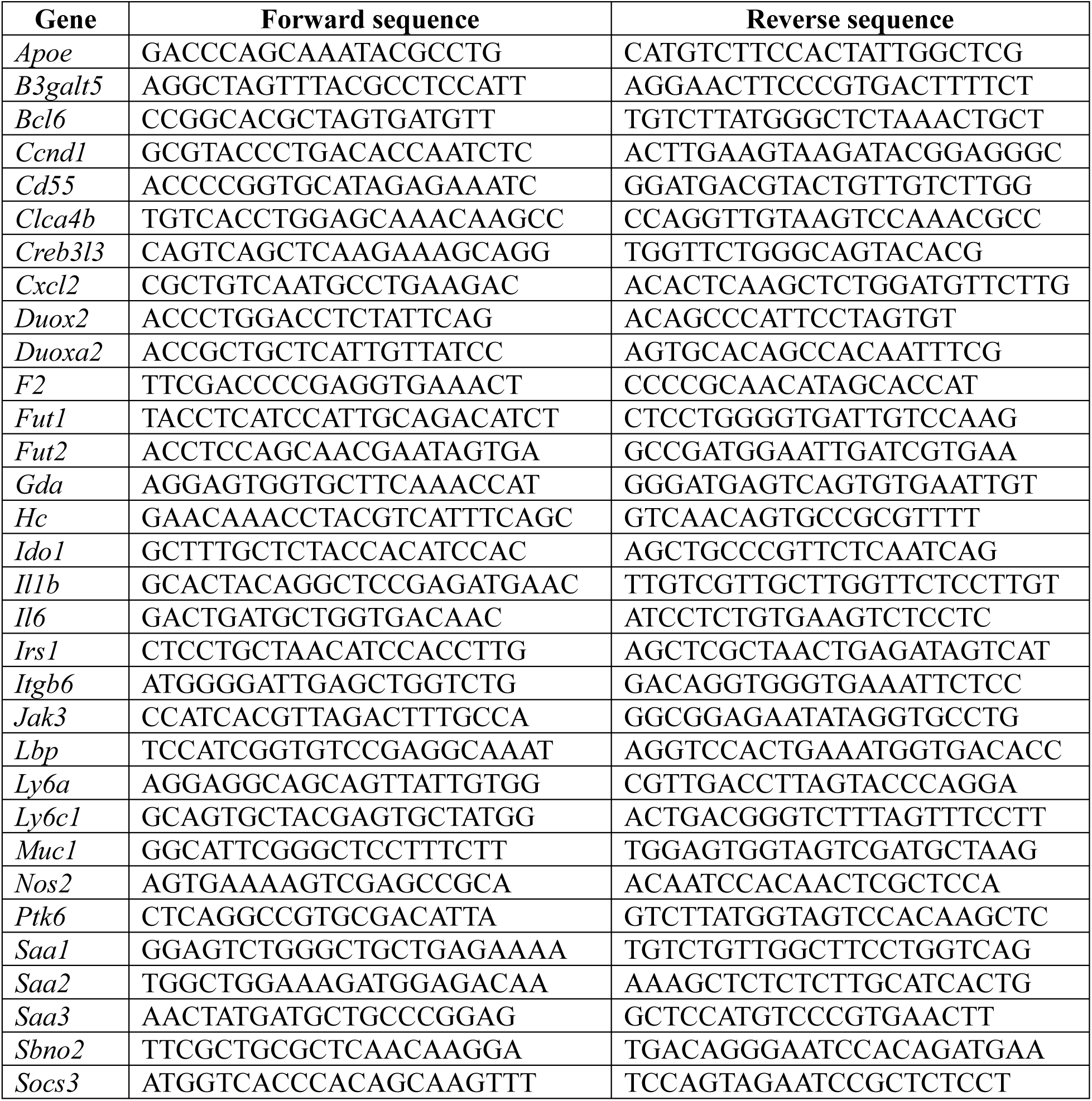
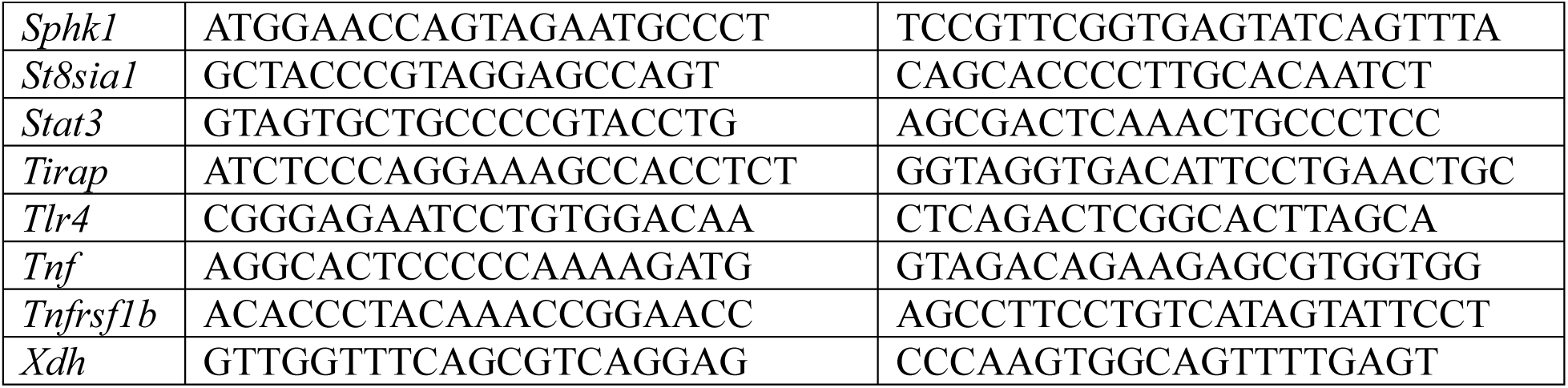
Primers (Forward and Reverse, Integrated DNA Technologies, Coralville, IA) used for Fluidigm analysis.

**Figure S1.**
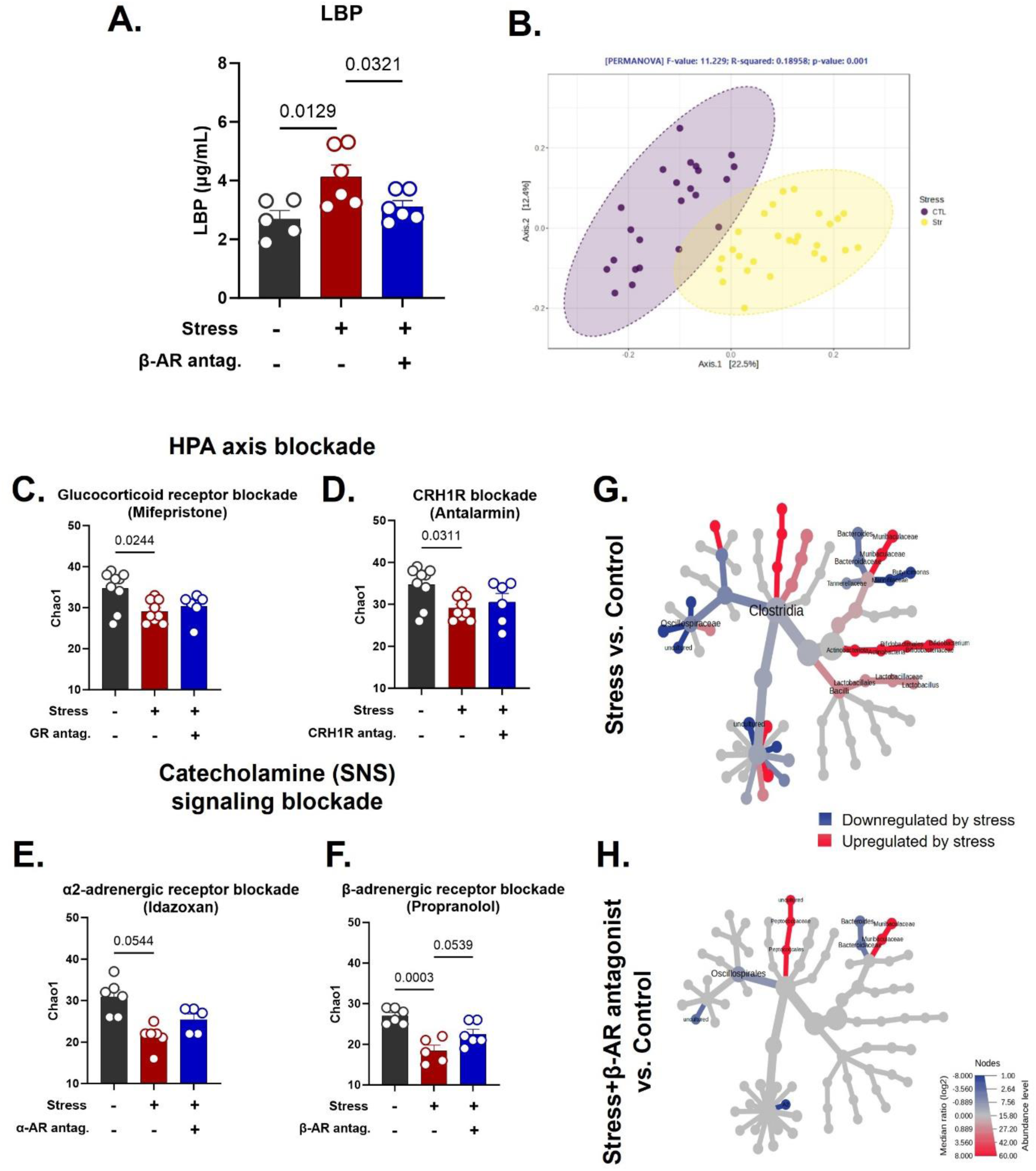
β-adrenergic receptor blockade prevents stress-induced increases in serum LBP and partially attenuates microbiome shifts associated with epithelial ROS/RNS responses. **(A)** Serum levels of lipopolysaccharide-binding protein (LBP) were measured by ELISA in mice exposed to social defeat stress (SDR) or left undisturbed, with or without treatment with the β-adrenergic receptor antagonist propranolol. Data were analyzed using one-way ANOVA followed by Tukey’s post hoc test. Statistical significance was set at *p* < 0.05. Values are expressed as mean ± SEM. n=5-6/group. **(B)** Principal coordinates analysis (PCoA) of Bray–Curtis dissimilarity reveals SDR-induced shifts in colonic microbiome beta-diversity. **(C–F)** Alpha-diversity assessed by Chao1 index from SDR mice treated with stress hormone receptor antagonists: **(C)** glucocorticoid receptor (mifepristone), **(D)** corticotropin-releasing factor receptor (antalarmin), **(E)** α₂-adrenergic receptor (idazoxan), and (**F)** β-adrenergic receptor (propranolol). **(G-H)** Heat tree visualizations of differential taxonomic abundance in **(G)** SDR vs. control mice and **(H)** SDR mice treated with propranolol vs. saline.

**Figure S2.**
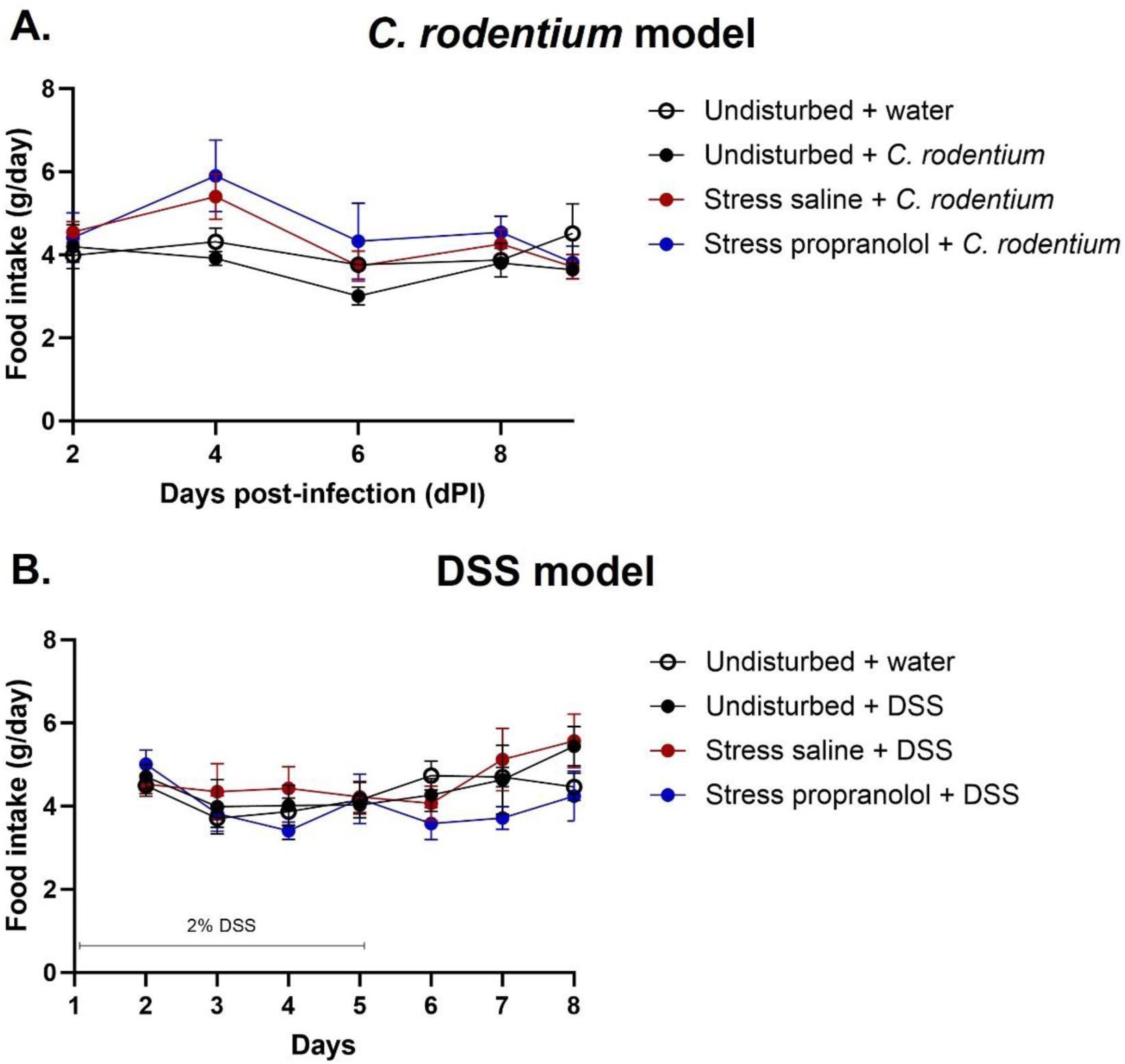
Food intake is not significantly altered by stress or propranolol treatment during *C. rodentium* infection or DSS-induced colitis. Food intake was monitored daily or every other day in two experimental models: (A) *Citrobacter rodentium* infection and **(B)** dextran sulfate sodium (DSS)- induced colitis. Data are presented as mean ± SEM (n = 5-6/group).

